# Hidden Contaminants in Sponge Genomes: Large-Scale Decontamination of 30 Public Assemblies

**DOI:** 10.64898/2026.06.12.731869

**Authors:** Kristian Bodulić, Kristian Vlahoviček

## Abstract

Sponges (phylum Porifera) are early-diverging metazoans that play central ecological roles and serve as models for understanding animal evolution. However, their associations with diverse microbial communities increase the risk of contamination in publicly available datasets, potentially compromising downstream biological inference. Despite growing genomic resources, systematic assessments of contamination in sponge genome assemblies have been lacking.

Here, we present a comprehensive contamination analysis of 30 publicly available sponge genome assemblies and introduce a reproducible and easily adoptable decontamination pipeline tailored to non-model organisms. Using this framework, we provide decontaminated versions of the analysed assemblies. The pipeline integrates three complementary lines of evidence: compositional outlier detection based on k-mer profiles and GC content, protein-level taxonomic classification using DIAMOND, and nucleotide-level classification with Kraken2. Scaffolds are designated as contaminants when supported by at least two independent signals. Pipeline performance was validated using a realistic spike-in dataset composed of bona fide sponge sequences and representative contaminant genomes.

The decontamination pipeline achieved 96.8% accuracy, 99.6% precision, and 90.8% recall, maintaining consistently strong recall across the vast majority of analyzed taxa. In addition, taxonomic assignments were accurately resolved to the genus level for 96.3% of identified contaminants. Application to public assemblies revealed variable contamination. On average, 14.5% of scaffolds per assembly were classified as contaminants, although they represented a low fraction of the total genome length, indicating that contamination is concentrated in relatively short scaffolds. Detected contaminants were dominated by bacterial phyla commonly associated with sponge microbiomes, including Pseudomonadota, Chloroflexota, and Poribacteria, with additional archaeal, protozoan, algal, and fungal sequences. Importantly, the number of complete BUSCO orthologs remained virtually unchanged following contamination removal, indicating minimal loss of genuine host scaffolds.

Taken together, our study provides 30 curated sponge genome assemblies and a consensus-based decontamination framework tailored to non-model organisms, improving the reliability of genomic resources for evolutionary, ecological, and functional analyses.

## Background

Sponges (phylum Porifera) are among the most ancient metazoan lineages and play fundamental ecological roles in marine and freshwater ecosystems [1–4]. Because they diverged early in animal evolution, sponges provide insights into the origins of multicellularity, the evolution of animal gene regulatory networks, and the emergence of key biological traits such as cell differentiation [1,2]. As sessile filter feeders, they mediate nutrient cycling, influence microbial community structure, and host diverse symbiotic assemblages collectively referred to as the sponge holobiont [3–8]. These associated microorganisms contribute to host nutrition, defense, and metabolite production, highlighting the evolutionary and ecological significance of sponge-microbe interactions [5–8].

The relationship between sponges and their microbiota presents unique opportunities for studying host-microbe coevolution, while also creating substantial challenges for genome assembly and downstream analyses [5–10]. Sponge tissues contain complex mixtures of bacterial, archaeal, eukaryotic, and viral material, with symbionts occurring at high abundance or possessing genomic features similar to those of the host [5–8]. Consequently, accurately distinguishing host-derived sequences from co-extracted microbial DNA is important for ensuring reliable genome assembly and annotation.

As genomic resources rapidly expand, assemblies deposited in public repositories increasingly serve as foundational datasets for comparative genomics, phylogenomics, and functional analyses. However, contamination remains a persistent issue in publicly available genome assemblies [11–14]. Contaminating material can originate from environmental DNA, symbiotic organisms, or laboratory sources, and its presence may bias gene prediction, distort functional annotations, and ultimately lead to incorrect biological inferences [11,12]. As such, accurate removal of contamination is essential for producing reliable reference genomes. Although several large-scale efforts have sought to mitigate contamination across databases such as RefSeq, these studies have largely overlooked sponge genomes, with limited attention given only to the original genome assembly of the model sponge *Amphimedon queenslandica* [12–14].

Several approaches have been developed to address contamination in genome assemblies, including composition-based methods, coverage-based clustering, marker gene detection, and taxonomic classification involving protein- or nucleotide-based similarity [12–19]. Although effective in many contexts, these methods are often applied as single-strategy frameworks or do not integrate multiple complementary signals into a unified scoring scheme [12–18]. Furthermore, some approaches operate exclusively on specific sequencing data types [17,18], and several tools are tailored to particular taxa, limiting their applicability on sponge genomes [18,19]. The performance of decontamination tools has also not been thoroughly evaluated in systems characterized by extensive horizontal gene transfer (HGT) or complex host-microbe associations, such as sponges, where distinguishing true host sequences from contamination is particularly challenging [5,6,8,12–19]. This difficulty is exacerbated by the limited representation of sponge and sponge-associated microbial sequences in reference databases, which constrains the accuracy of similarity-based classification.

Here, we present a large-scale contamination assessment of publicly available sponge genome assemblies. Our study delivers a curated set of 30 decontaminated genomes, a taxonomic characterization of contaminants, and a reproducible decontamination pipeline designed for non-model organisms. By revealing the extent and diversity of previously hidden contaminants in public sponge assemblies, this work provides improved genomic resources, while highlighting the importance of contamination-aware analyses in holobiont systems.

## Methods

### Genome Decontamination Pipeline

The genome decontamination pipeline developed in this study integrates multiple lines of evidence to generate a unified contamination classification, assign taxonomic annotations to contaminant scaffolds, and produce a detailed contamination report (Figure 1). The pipeline takes as input a genome assembly in FASTA format together with the target taxon label. By default, only scaffolds longer than 1 kb are analyzed. Although the pipeline offers several tunable parameters, they were not optimized for the datasets analyzed in this study. Default parameter values were selected to reflect biologically reasonable settings, with their rationale provided in Supplementary Table 1.

**Figure 1.**
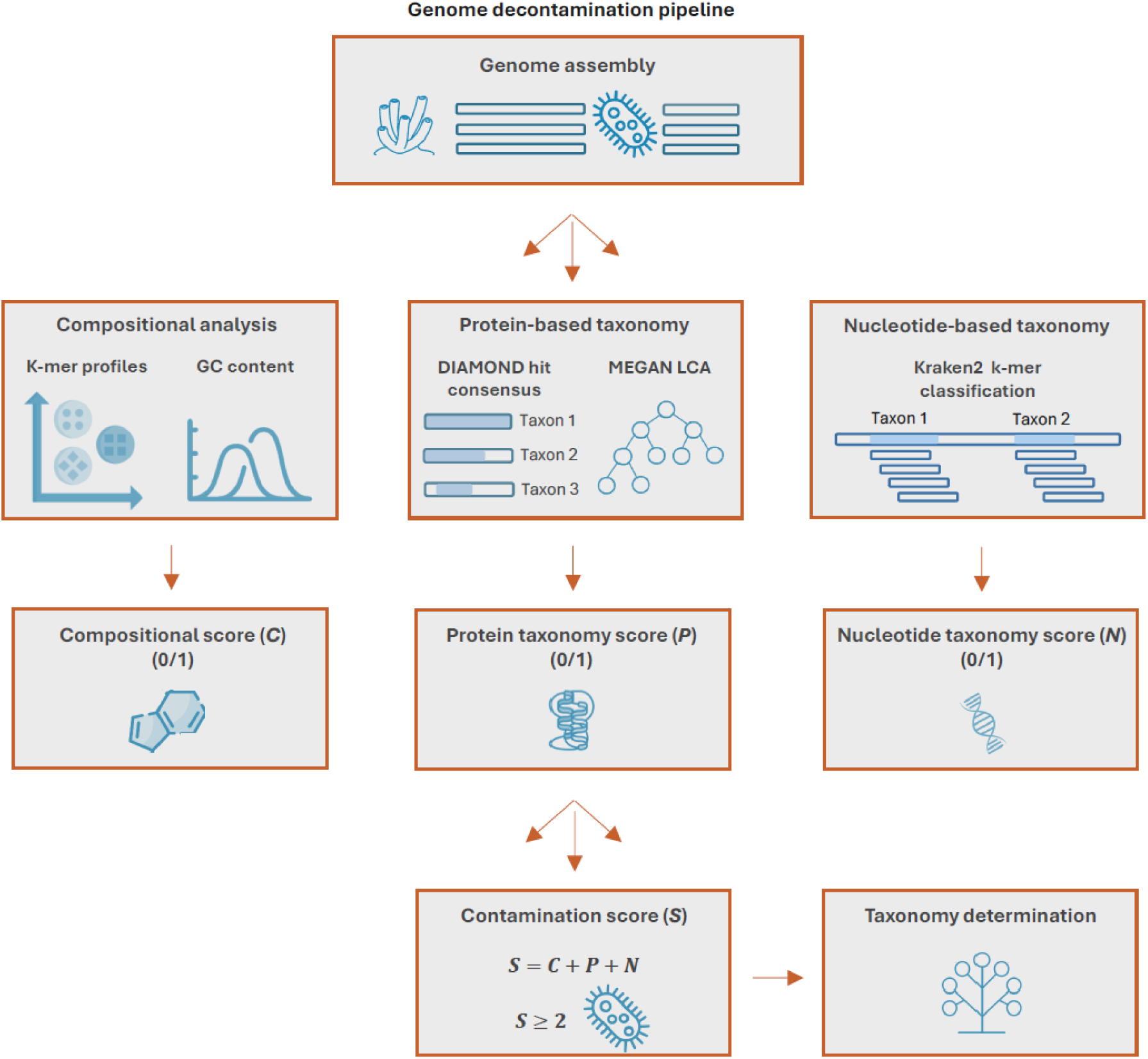
Overview of the genome decontamination pipeline. The pipeline takes a genome assembly as input and evaluates each scaffold using three independent lines of evidence. This includes (i) compositional outlier detection based on k-mer profiles and GC content (compositional score), (ii) protein-level taxonomic classification using DIAMOND hit consensus and MEGAN LCA (protein taxonomy score), and (iii) nucleotide-level taxonomic classification using Kraken2 (nucleotide taxonomy score). Binary scores are integrated into a final contamination score, with scaffolds classified as contaminants when at least two lines of evidence suggest contamination. Contaminant scaffolds are subsequently assigned taxonomic labels based on the available nucleotide- or protein-based evidence. **Abbreviations**: LCA: lowest common ancestor, *C*: compositional score, *P*: protein taxonomy score, *N*: nucleotide taxonomy score, *S*: contamination score.

Specifically, the decontamination pipeline computes three binary scores for each scaffold: a compositional score based on k-mer frequencies and GC content, a protein score derived from taxonomic assignments of encoded proteins, and a nucleotide score based on taxonomic k-mer classification. Scaffolds are classified as contaminants based on the integrated outcomes of these scoring components. Detailed descriptions of each scoring metric are provided below.

The compositional score identifies scaffolds with atypical k-mer frequency profiles or GC content. Scaffold sequences are decomposed into rolling k-mers of length 5, and each k-mer is collapsed with its reverse complement into a single canonical representation defined as the lexicographically smaller sequence (R package Biostrings) [20,21]. For each scaffold, canonical k-mer counts are converted to proportions and transformed using the centered log-ratio (CLR) to account for compositional effects inherent to relative abundance data (R package compositions) [22]. Principal component analysis (PCA) is performed on the CLR-transformed k-mer profiles, retaining the top principal components that together explain at least 90% of the transformed k-mer frequency variance, up to a maximum of 20 components.

The resulting representation is used for unsupervised clustering with Gaussian finite mixture models (R package mclust) [23]. Models are fitted for a range of cluster numbers ranging from 1 to a user-defined maximum (default = 10) and evaluated across several covariance structures (EII, EEI, EEE, VII, VEI, and VVI). The optimal model is selected using the integrated completed likelihood criterion. The reference cluster is defined as the cluster with the highest cumulative scaffold length, as these scaffolds are assumed to represent the target genome rather than contaminants. Mahalanobis distances between each cluster centroid and the reference cluster centroid are computed in the retained PCA space. Clusters are flagged as compositional outliers if the square root of this distance exceeded a heuristic chi-square distribution–based cutoff (α = 0.005 by default, degrees of freedom equal to the number of retained principal components). All scaffolds assigned to outlier clusters are flagged as compositional outliers.

Following k-mer decomposition, GC content is calculated for each scaffold, and deviations from the genome mean are quantified using z-scores. Scaffolds with absolute z-scores exceeding a threshold (default = 2.5) are flagged as GC outliers. The compositional score (*C*) is defined as a binary indicator equal to 1 if a scaffold is identified as an outlier based on either k-mer composition or GC content, and 0 if no compositional deviation is detected.

Protein-based detection of taxonomic outliers is performed using two complementary strategies: a custom DIAMOND consensus method and a lowest common ancestor (LCA) approach implemented in MEGAN. In the DIAMOND consensus method, each scaffold is searched against the NCBI non-redundant protein database using DIAMOND blastx with sensitive parameters optimized for genomic scaffolds (--sensitive, - e 1e-5, --range-culling, -F 15, --top 10) [24,25]. All subject hits are assigned taxonomic identifiers (R package taxonomizr) and classified into one of 14 kingdom-level taxonomic groups (Bacteria, Archaea, Viridiplantae, Haptophyta, Rhodophyta, Stramenopiles, Alveolata, Excavata, Rhizaria, Choanoflagellata, Amoebozoa, Fungi, Metazoa, and Viruses) [26]. Overlapping DIAMOND hits along genomic coordinates are collapsed into taxon-specific DIAMOND hit sets (R package GenomicRanges) [27]. Each hit set is assigned a weight *S_w_* defined as:

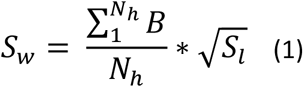

Here, *N_h_* represents the number of DIAMOND hits per hit set, *B* is the hit bitscore and *S_l_* marks the hit set length. This definition ensures that hit sets are weighted according to alignment quality and length.

Each scaffold-taxon combination is assigned a weight *W* given by:

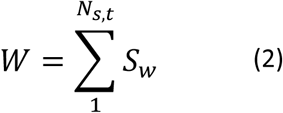

where *N_s_*_,*t*_ represents the number of hit sets assigned to taxon *t* on scaffold *s*.

Under the DIAMOND consensus framework, a scaffold is classified as an outlier if the maximum weight of any non-target taxon exceeds the target taxon weight multiplied by a specified factor (default = 1.2). Additionally, the total length of the highest-weight non-target taxon hit sets must exceed a minimum fraction of the scaffold length (default = 0.2) and must be greater than the total target taxon hit set length.

Independently, each scaffold is assigned a taxonomic label using MEGAN on the same DIAMOND results with parameters optimized for genomic scaffolds (-lg, -alg longReads, -me 1e-5) [28]. Scaffolds assigned to a non-target taxon via the LCA algorithm implemented in MEGAN are considered outliers. The protein taxonomy score (*P)* is defined as a binary indicator equal to 1 if a scaffold is identified as an outlier by either the DIAMOND consensus or MEGAN LCA approach, and 0 if no evidence of taxonomic inconsistency is detected.

Nucleotide-based detection of taxonomic outliers is performed using Kraken2, with scaffolds queried against the NCBI core nucleotide database using default parameters [25,29]. Kraken2 assigns scaffold taxonomy by classifying overlapping k-mers based on exact matches to database k-mers. All classified k-mers are mapped to taxonomic identifiers and aggregated into the same set of 14 taxonomic groups used in the protein-based analysis.

For each scaffold–taxon combination, an absolute k-mer fraction *A* is calculated as:

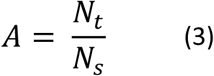

where *N_t_* denotes the number of k-mers assigned to a given taxon and *N_s_* represents the total number of k-mers per scaffold.

A scaffold is classified as a nucleotide-based taxonomic outlier if both of the following conditions are satisfied: (i) the maximum absolute k-mer fraction among non-target taxa exceeds the target taxon k-mer fraction multiplied by a specified factor (default = 1.2) and (ii) the maximum non-target k-mer fraction exceeds an absolute threshold (default = 0.1).

The nucleotide taxonomy score (*N*) is defined as a binary indicator, taking the value 1 when a scaffold is identified as a nucleotide-based taxonomic outlier and 0 when no evidence of taxonomic discrepancy is detected.

An overall scaffold contamination score (*S*) is defined to integrate evidence from compositional, protein, and nucleotide scoring components. For each scaffold, the score is calculated as:

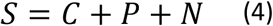

A scaffold is classified as a contaminant when *S* >= 2, indicating support for contamination from at least two independent lines of evidence.

Taxonomic labels are assigned to scaffolds identified as contaminants at the highest possible resolution using evidence from nucleotide- or protein-based analyses. In both cases, the maximum considered taxonomic resolution is the genus level. When nucleotide-based detection indicates a taxonomic signal (*N* = 1), scaffolds are assigned to the non-target taxon with the highest number of matching k-mers within a given taxonomic rank. Taxonomic ranks are evaluated hierarchically, beginning at the genus level and proceeding to higher ranks until the following criteria are satisfied: (i) the absolute k-mer fraction for the taxon exceeds a minimum threshold (default = 0.1) and (ii) the relative k-mer fraction for the same taxon exceeds a second threshold (default = 0.5). The relative k-mer fraction *R* is defined as:

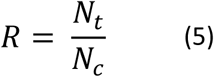

where *N_c_* denotes the total number of taxonomically classified k-mers per scaffold.

If nucleotide-based evidence is absent (*N* = 0), or if the above criteria are not satisfied at any taxonomic rank, contaminant scaffolds are assigned to the most specific rank supported by protein alignments. Specifically, the assigned label corresponds to the taxon with the greatest cumulative length of DIAMOND hit sets associated with the scaffold. Taxonomic ranks are evaluated hierarchically until the following criteria are satisfied: (i) the cumulative length of hit sets assigned to the taxon exceeds a minimum fraction of the scaffold length (default = 0.25) and (ii) exceeds a minimum fraction of the total hit set length per scaffold (default = 0.5). If these criteria are not met for any taxonomic rank, the scaffold is assigned an unknown taxonomy. This hierarchical procedure prioritizes nucleotide-based assignments when available, while leveraging protein-based evidence to provide taxonomic resolution for scaffolds lacking sufficient nucleotide signal.

### Genome Decontamination Pipeline Validation

To evaluate pipeline performance, a spike-in validation dataset was constructed comprising bona fide sponge genomic sequences alongside a representative set of contaminant genomes. Sponge sequences were derived from chromosome-level genome assemblies spanning 18 species. For each genome, only chromosomal sequences were retained, while unlocalized scaffolds were excluded based on genome annotation. In addition, five mitochondrial sequences were included to supplement the sponge dataset. A diverse panel of contaminant taxa was incorporated to reflect common sources of non-target sequences in aquatic ecosystems and sponge microbiomes, encompassing typical symbionts as well as environmental organisms across a broad taxonomic range. Specifically, the dataset included genome assemblies from 71 species representing bacteria, archaea, protozoa, algae, fungi, and viruses, along with five bacterial plasmid sequences. Detailed information on genome assemblies included in the validation dataset is provided in Supplementary Table 2 (sponges) and Supplementary Table 3 (contaminants).

The validation dataset was constructed by fragmenting the source sequences to reproduce scaffold length distributions observed in the sponge assemblies analysed for contamination. Only sequences longer than 1 kb were considered, as shorter sequences were not classified by the decontamination pipeline. The length distribution of sponge scaffolds analysed for contamination was quantified using 13 predefined length bins (1–2 kb, 2–4 kb, 4–10 kb, 10–20 kb, 20–30 kb, 30–50 kb, 50–100 kb, 100–200 kb, 200–500 kb, 500 kb–1 Mb, 1–5 Mb, 5–10 Mb, and 10–100 Mb). The proportion of scaffolds within each bin was used to guide sampling of fragments from the source validation sequences to match empirical assembly profiles. Fragments were generated by sampling contiguous segments from source sequences, with fragment lengths randomly drawn from length bins. Fragment coordinates were sampled uniformly along each source sequence. For sponge source sequences, remaining segments longer than 1 kb were returned to the sampling pool, with reuse permitted up to four times per source sequence. Fragments containing more than 10% ambiguous bases were excluded from the final sequence set.

The resulting validation dataset comprised 13834 sequences, of which 4653 (33.6%) originated from contaminants. The N_50_ value of sponge-derived sequences was 7.01 Mb, while contaminant-derived sequences exhibited an N_50_ of 1.68 Mb. Comparison between sequence lengths in the validation dataset and sponge genome assemblies analysed for contamination is shown in Supplementary Figure 1. The decontamination pipeline was run on the validation dataset using default parameters, except for the maximum number of k-mer–based scaffold clusters. The decontamination pipeline was applied to the validation dataset using default parameters, with the exception of the maximum number of k-mer-based scaffold clusters. This parameter was increased to 20 to accommodate the greater species diversity present in the validation dataset, which exceeds that typically expected in a single analyzed genome. The following software and database versions were used: DIAMOND v2.1.13, NCBI non-redundant database downloaded on September 17, 2025, MEGAN v7.1.1, MEGAN taxonomy mapping file v7.0.0, Kraken2 v2.1.6, NCBI core nucleotide database downloaded on October 29, 2025, taxonomizr v0.11.1, taxonomizr database downloaded on December 27, 2025, Biostrings v2.76.0, compositions v2.0.9, mclust v6.1.2, and GenomicRanges v1.62.1. Validation dataset generation and result analysis were performed using custom R scripts (R v4.5.2), with the following R packages used for data manipulation and visualization: data.table (v1.17.8) [30], ggplot2 (v4.0.2) [31], ComplexUpset (v1.3.3) [32], and pheatmap (v1.0.13) [33].

### Sponge Genome Assembly Decontamination

The genome decontamination pipeline was applied to 30 contiguous sponge genome assemblies retrieved via the NCBI Genomes resource, including assemblies from GenBank (*n* = 29) and RefSeq (*n* = 1, *Sycon ciliatum*) [25]. The dataset was primarily composed by Demospongiae (*n* = 27), with additional representatives from Calcarea (*n* = 1), Hexactinellida (*n* = 1), and Homoscleromorpha (*n* = 1). The complete list of analyzed assemblies is provided in Supplementary Table 4. All assemblies were processed using the decontamination pipeline with default parameters, with software dependencies and database versions identical to those used in the validation dataset analysis. The resulting outputs were analyzed using a custom R script, with the following R packages used for data manipulation and visualization: data.table (v1.17.8), ggplot2 (v4.0.2), pheatmap (v1.0.13), and ggVennDiagram (v 1.5.4) [34].

### Assessment of Genome Completeness Using BUSCO

To evaluate potential false-positive contaminant classification, we compared genome completeness between the original and decontaminated sponge genome assemblies. Completeness was assessed using BUSCO (v6.0.0) with the Metazoa lineage dataset [35]. The resulting metrics were processed and visualized using a custom R script, with R packages data.table (v1.17.8) and ggplot2 (v4.0.2).

## Results

### Genome Decontamination Pipeline Validation

To systematically assess and remove contamination in sponge genomes, we developed a dedicated decontamination pipeline and applied it to 30 publicly available assemblies. This effort generated three main outcomes: a curated set of decontaminated sponge genomes, a broadly applicable decontamination workflow, and a detailed characterization of contamination patterns across assemblies. Our approach combines complementary lines of evidence to improve classification accuracy. Specifically, the decontamination pipeline integrates a compositional score that assesses k-mer profiles and GC content, a protein score derived from taxonomic classification of encoded proteins, and a nucleotide score based on k-mer taxonomic profiling.

We first validated the pipeline using a validation spike-in dataset comprising bona fide sponge sequences alongside a diverse set of representative contaminants (Figure 2). The pipeline demonstrated high overall accuracy in classifying contaminant scaffolds (96.8%), with strong precision (99.6%) and recall (90.8%) (Figure 2a). Among the individual components, the compositional score showed the weakest performance (accuracy 74.6%, precision 59.5%, recall 77.1%), whereas protein and nucleotide taxonomy scores exhibited stronger and comparable performance (protein: accuracy 96.2%, precision 98.2%, recall 90.5%; nucleotide: accuracy 96.0%, precision 99.4%, recall 88.8%). The integrated pipeline achieved the highest overall accuracy, outperforming each individual component, although performance gains relative to the protein and nucleotide scores were modest. All sponge mitochondrial sequences were correctly classified despite their prokaryotic features, highlighting low false-positive rates for atypical host sequences. Additionally, only one (6.7%) bacterial plasmid sequence was falsely classified as non-contaminant, demonstrating the pipeline’s ability to accurately classify extrachromosomal sequences.

**Figure 2.**
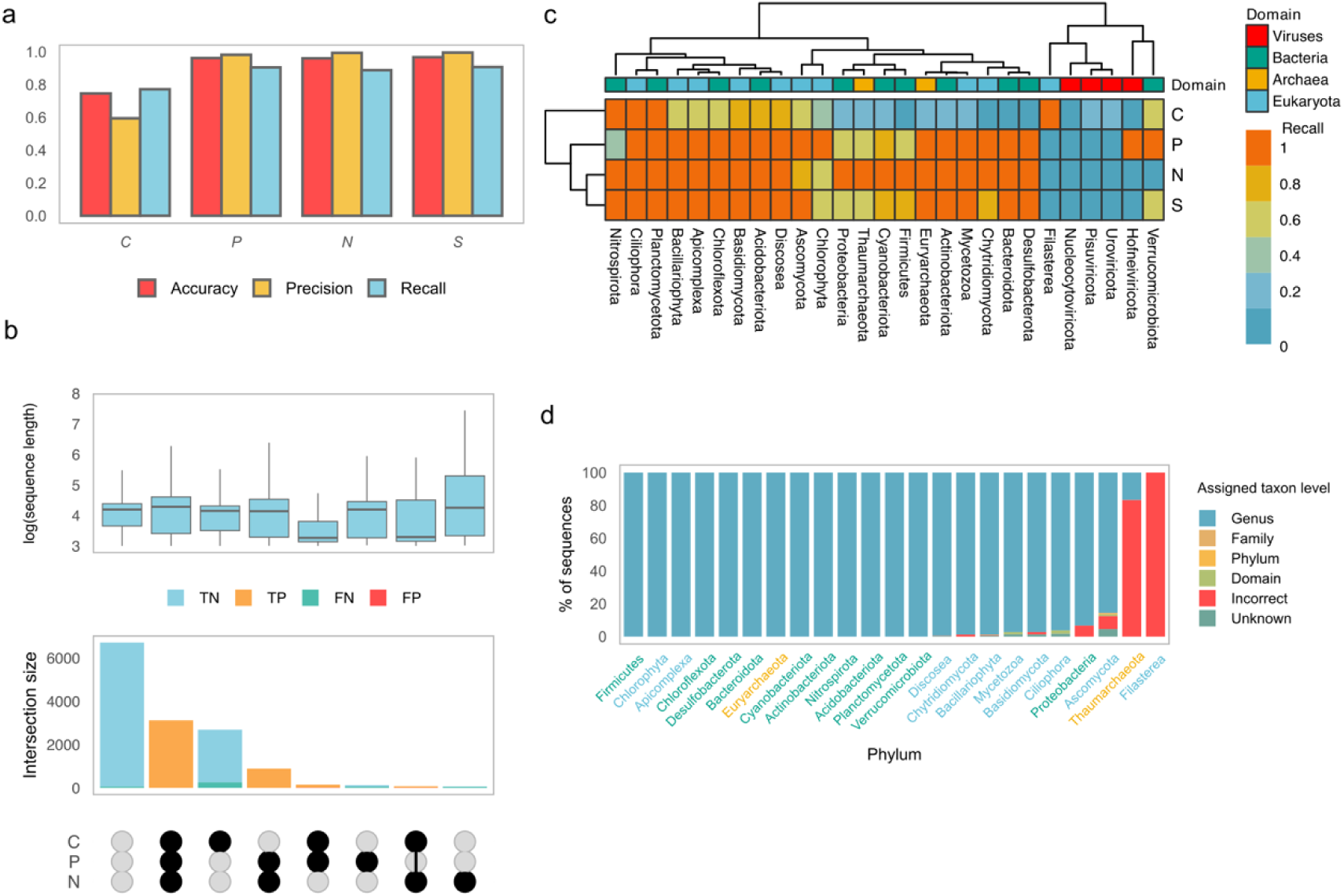
Validation spike-in dataset results. Performance of the decontamination pipeline on a realistic spike-in dataset composed of bona fide sponge sequences and representative contaminants. **a** Accuracy, precision, and recall for individual scoring components and for the integrated contamination score. **b** Intersection analysis showing agreement among scoring components. Bar heights indicate the number of sequences in each intersection, while boxplots summarize sequence length distributions for each intersection. Boxplots represent the median and IQR, with whiskers extending to ±1.5 × IQR. Colors indicate classification results. **c** Heatmap showing recall across contaminant phyla for scoring components and for the final integrated classification, with hierarchical clustering applied to phyla and scoring methods. Contaminant phyla are colored by taxonomic domains. **d** Accuracy of taxonomic assignment for contaminant sequences, showing the proportion of sequences resolved to different taxonomic ranks as well as incorrect and unknown classifications. X-axis labels represent contaminant phyla and are colored by taxonomic domains analogous to panel c. **Abbreviations**: *C*: compositional score, *P*: protein taxonomy score, *N*: nucleotide taxonomy score, *S*: contamination score, TN: true negatives, TP: true positives, FN: false negatives, FP: false positives.

The intersection analysis revealed strong agreement among the scoring components (Figure 2b). The majority of non-contaminant sequences were correctly classified by all three components (6649, 72.4% of true non-contaminants). Similarly, 3120 sequences (67.1% of true contaminants) were correctly classified as contaminants by all three scores, while 891 sequences (19.2% of true contaminants) were correctly classified as contaminants by protein and nucleotide scores, but not by the compositional score. In 141 cases (3.0% of true contaminants), the compositional and protein components correctly identified sequences as contaminants, whereas the nucleotide component failed to detect them. Similarly, 71 sequences (1.5% of true contaminants) were correctly classified as contaminants by the compositional and nucleotide scores but missed by the protein component. Among scaffolds supported by a single signal, the compositional score alone falsely classified 2430 sequences as contaminants (26.5% of true non-contaminants), indicating a higher false positive rate when used in isolation. However, 257 contaminant sequences (5.5% of true contaminants) were detected by the compositional score but falsely classified as non-contaminants by both protein and nucleotide components.

Overall, the length distribution across intersections did not suggest a strong length-associated bias. An exception was observed for scaffolds missed by the nucleotide score but detected by both compositional and protein components, which tended to be notably short (median 1.79 kb, IQR 1.33–4.35 kb).

Recall varied across taxonomic groups and scoring methods, with modest differences observed among taxonomic domains (Figure 2c). The integrated pipeline achieved consistently high recall across most phyla, a pattern that was also observed for the protein and nucleotide taxonomy scores. In contrast, the compositional score showed lower recall in several prokaryotic and eukaryotic phyla. Notably the compositional score successfully detected contaminants from the Filasterea phylum that were not captured by the taxonomy-based methods. Viral phyla generally exhibited low recall across all components and the final integrated score.

We also evaluated the accuracy of taxonomic assignments for contaminant sequences (Figure 2d). A taxonomic assignment was considered correct if the assigned lineage matched any taxonomic rank along the true lineage of the taxon. Using this definition, 4223 (97.7%) contaminant sequences were correctly classified overall, with 4068 (96.3%) sequences accurately resolved at the genus level. Only 63 (1.5%) sequences were classified as unknown and were not assigned any taxonomy. The pipeline exhibited strong performance across the vast majority of contaminants, with only two phyla showing low accuracy (Filasterea; 0%, Thaumarchaeota; 16.7%). Interestingly, all misclassified Thaumarchaeota sequences were assigned to alternative archaeal genera, whereas Filasterea sequences were misclassified as fungal or algal taxa. These results indicate generally robust taxonomy assignment with limited taxon-specific variation.

### Sponge Genome Assembly Decontamination

We next applied the decontamination pipeline to 30 publicly available sponge genomes assemblies (Figure 3, Supplementary Table 5). The analysis revealed variable levels of contamination across datasets, as reflected by the number of scaffolds classified as contaminants (Figure 3a). The mean proportion of contaminant scaffolds was 14.5% (SD 16.6%), with only *Corticium candelabrum* and *Petrosia crassa* showing no detectable contamination. The highest proportions of contaminant scaffolds were observed in *Xestospongia muta* (200, 66.9%), *Rhabdastrella globostellata* (156, 47.4%), *Spongilla officinalis* (40, 37.7%), *Axinella damicornis* (239, 36.0%), *Diacarnus erythraeanus* (50, 34.3%), and *Phyllospongia foliascens* (562, 31.1%).

**Figure 3.**
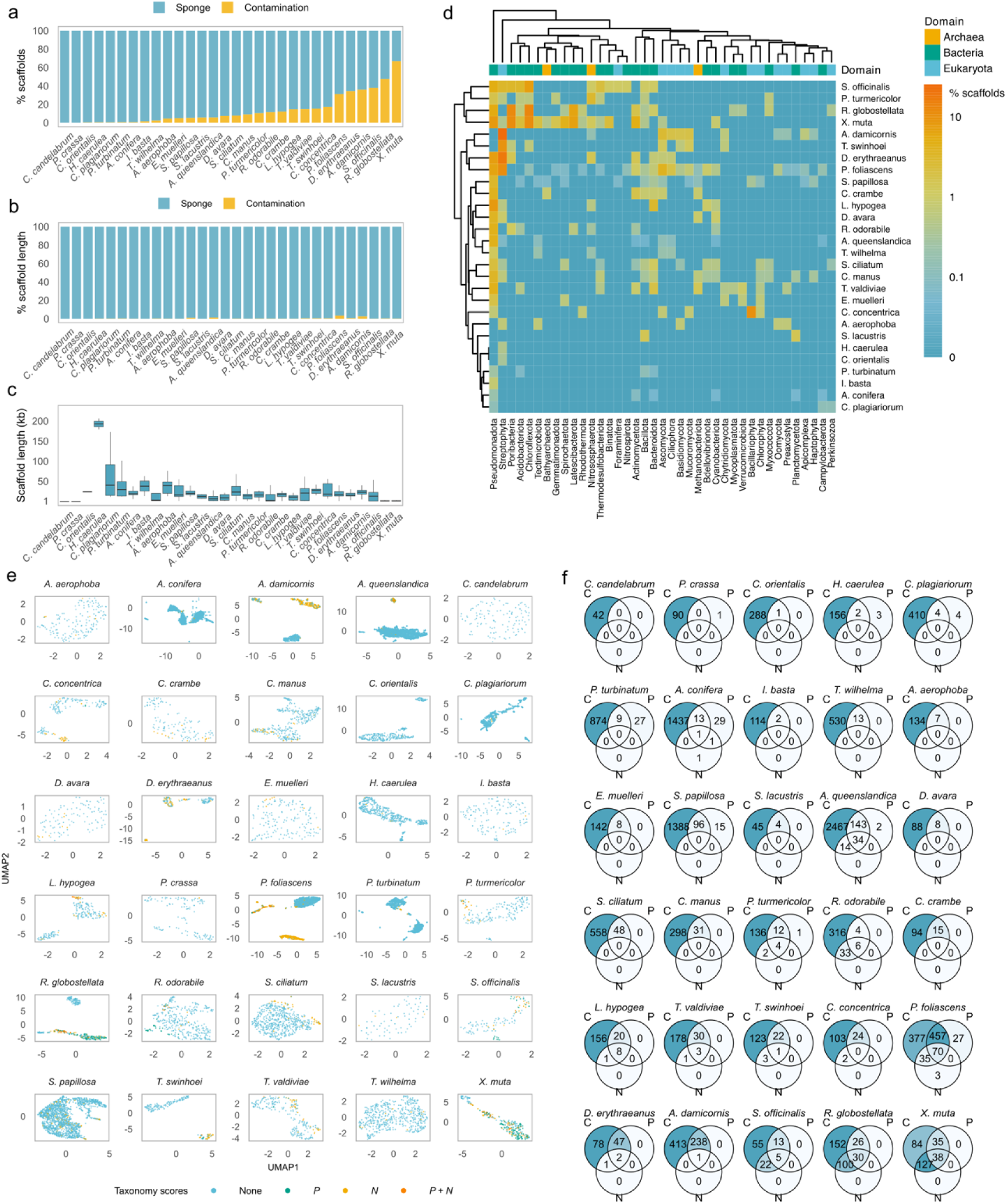
Decontamination analysis of 30 publicly available sponge genome assemblies. Application of the genome decontamination pipeline to scaffold-level sponge assemblies retrieved from public databases. **a** Proportion of scaffolds classified as sponge-derived or contaminants for each assembly. **b** Proportion of total genome length attributed to sponge-derived and contaminant scaffolds. **c** Length distribution of contaminant scaffolds across assemblies. Boxplots represent the median and IQR, with whiskers extending to ±1.5 × IQR **d** Heatmap showing the proportion of contaminant scaffolds across sponge assemblies, with hierarchical clustering applied to contaminant phyla and sponge species. Contaminant phyla are colored by taxonomic domains. **e** Intersection analysis illustrating agreement among the compositional, protein, and nucleotide scoring components for each genome assembly. **f** UMAP projection of principal components derived from k-mer profiles used in compositional scoring, highlighting scaffolds supported by different combinations of taxonomic scores. **Abbreviations**: *C*: compositional score, *P*: protein taxonomy score, *N*: nucleotide taxonomy score, *S*: contamination score.

Contamination levels were lower when expressed as the percentage of total genome length, with a mean of 0.5% (SD 0.7%). The highest proportions of contaminated genome length were observed in *P*. *foliascens* (3.5%) and *A*. *damicornis* (2.5%). The stated assemblies also exhibited a high number of contaminant scaffolds. Across all species, the median contaminant scaffold length was 15.0 kb (range 1.0–228.5 kb) (Figure 3c). Several species with a high number of contaminant scaffolds showed relatively low contaminant scaffold lengths, including *X*. *muta* (median 1.0 kb, range 1.0–25.7 kb) and *R*. *globostellata* (median 1.0 kb, range 1.0–36.8 kb).

Next, we performed taxonomic classification of the detected contaminant scaffolds (Figure 3d). Bacterial phyla generally dominated the contaminant profiles. Pseudomonadota was the most abundant bacterial phyla, with 467 scaffolds identified across 22 sponge species. This was followed by Chloroflexota (77 scaffolds detected in six species), Poribacteria (74 scaffolds in six species), and Bacteroidota (45 scaffolds distributed across 12 species). Furthermore, four archaeal phyla were detected. The most prevalent were Nitrososphaerota, with 15 scaffolds identified in eight species, and Methanobacteriota, with 11 scaffolds found in six species. Among eukaryotic contaminants, algal sequences were particularly abundant, primarily belonging to green algae (Streptophyta, 420 scaffolds detected in 15 species). Fungal contaminants were also common and were mainly assigned to Ascomycota (101 scaffolds in 10 species).

The greatest contaminant diversity was observed in *P*. *foliascens*, which harbored 23 distinct phyla. The most abundant were Streptophyta (195 scaffolds, 11.1%), Pseudomonadota (111 scaffolds, 6.3%), and Ascomycota (69 scaffolds, 3.9%). High contaminant diversity was also detected in *X*. *muta*, with 18 identified contaminant phyla, 14 of which were bacterial. The most prevalent phyla were Poribacteria (29 scaffolds, 11.5%), Chloroflexota (28 scaffolds, 11.2%), Pseudomonadota (21 scaffolds, 8.4%), and Latescibacterota (18 scaffolds, 7.2%). Similarly, *R*. *globostellata* exhibited substantial contaminant diversity, with 14 phyla detected, 12 of which were bacterial. The dominant groups were Chloroflexota (38 scaffolds, 12.6%), Poribacteria (35 scaffolds, 11.7%), and Latescibacterota (19 scaffolds, 6.3%).

We next assessed concordance among the three contamination-scoring components for each sponge assembly. In contrast to the validation dataset, the majority of contaminant scaffolds across all sponges were supported by two of the three scoring components (Figure 3e). Only 203 scaffolds (10.9%) were classified as contaminants by all three scores. The largest fraction of contaminant scaffolds (1332, 70.8%) was identified exclusively by the compositional and protein scores, whereas 341 contaminant scaffolds (18.2%) were supported by the compositional and nucleotide scores. Consistent with observations from the validation dataset, the compositional score most frequently flagged scaffolds as contaminants and would likely yield a substantial number of false positives if applied in isolation.

Agreement between the compositional and taxonomy-based scoring components is further illustrated by a UMAP projection of principal components derived from the k-mer profiles used for compositional scoring (Figure 3f). Scaffolds classified as contaminants by both protein and nucleotide taxonomy scores generally formed clusters distinct from scaffolds lacking taxonomic evidence of contamination. This separation was particularly pronounced in assemblies exhibiting high levels of contamination. Collectively, these results underscore the importance of the integrated scoring framework implemented in our pipeline.

Finally, we evaluated the potential for false-positive contaminant detection by comparing the number of complete BUSCO orthologs (Metazoa dataset) before and after genome decontamination (Supplementary Figure 2). Notably, 25 assemblies (83.3%) showed identical completeness before and after decontamination. The remaining five assemblies exhibited only a minimal reduction in completeness, with the loss of a single BUSCO ortholog. These assemblies included *Crambe crambe*, *Halisarca caerulea*, *Penares turmericolor*, *P*. *foliascens*, and *Sycon ciliatum*.

## Discussion

The rapid expansion of genomic resources for non-model metazoans has advanced our understanding of early animal evolution, while also introducing a significant challenge regarding data integrity. Sponges present a unique bottleneck for genomic assembly due to the close association between host cells and a diverse microbial community [5–8]. In this study, we developed an integrated decontamination pipeline and applied it to 30 publicly available sponge assemblies. While the total percentage of contaminant genome length remains relatively low, our results suggest a substantial number of contaminating scaffolds. Notably, several of the analysed assemblies had reached chromosome-level resolution, yet contamination was still detected, predominantly within short unlocalized scaffolds. These non-target sequences potentially harbor thousands of misannotated genes that could bias downstream evolutionary and functional analyses. Importantly, we provide systematically decontaminated versions of the analysed sponge genome assemblies, offering a more reliable resource for future genomic studies.

Validation using a spike-in dataset demonstrated strong decontamination pipeline performance, with an overall accuracy of 96.8%, a precision of 99.6%, and a recall of 90.8%. These metrics suggest that the integrated scoring system is highly reliable for classifying host and contaminant scaffolds. However, the high recall observed in the validation dataset is likely an optimistic reflection of real-world performance considering that the included source sequences are represented in the databases utilized by the pipeline. The signal from similarity-based methods may be weaker when processing novel assemblies. Consequently, decontamination efficacy is inherently constrained by the phylogenetic breadth of existing biological catalogs. As genomic efforts move toward increasingly obscure lineages, the importance of the compositional score increases, as it remains the only line of evidence independent of external database representation.

It should be noted that the validation dataset did not fully mimic typical genome assemblies, as it comprised fragments from multiple sponge species alongside a wide range of contaminants. While this strategy potentially introduced bias in compositional scoring, it represented a strategic design choice to enable a broader characterization of pipeline performance across a diverse taxonomic range. Notably, this multi-genome composition created a more rigorous test for compositional scoring than a typical genome assembly, as the pipeline had to navigate a significantly higher level of genomic heterogeneity to accurately cluster both target and contaminant sequences.

The BUSCO results provide another validation of our decontamination methodology. The near-identical completeness observed in original and decontaminated sponge genomes goes in hand with high precision demonstrated in the validation dataset. These results indicate that the pipeline successfully removed non-target DNA without compromising the integrity of the host genome. The loss of a single BUSCO ortholog in five analysed assemblies may be attributable to the removal of short scaffolds (<1 kb), which were automatically excluded by the decontamination pipeline.

An important observation in this study was the significant divergence in scoring component concordance between the validation dataset and the public sponge assemblies. In the controlled environment, the majority of contaminant sequences were flagged by all three scores (*C*, *P*, and *N*). In contrast, contaminants identified within the public sponge assemblies were predominantly supported by two components, underscoring the necessity of the integrated scoring framework. Relying on a single line of evidence would yield high false-positive rates, while requiring a triple-hit consensus would result in poor recall. This may be a direct consequence of the underrepresentation of sponge-associated microbial sequences in public repositories, necessitating a multi-modal approach to ensure robust detection.

A core design choice of our pipeline is the aggregation of DIAMOND and Kraken2 hits into 14 kingdom-level taxonomic bins. This strategy addresses the substantial underrepresentation of sponge sequences in public databases. Genes originating from non-model animals frequently return their strongest matches to other metazoans, whose genomes are more comprehensively represented. By collapsing all animal hits into a single metazoan bin, the pipeline minimizes the risk of falsely classifying host-derived scaffolds as contaminants. However, this design introduces an important limitation: taxonomy-based scoring components cannot distinguish between organisms within the same kingdom. Consequently, the pipeline is unable to detect intra-kingdom contamination, such as human DNA introduced during laboratory handling or sequences originating from other marine metazoans. In practice, we consider this limitation acceptable, as intra-kingdom contaminants are likely far less prevalent than ecological contaminants, which the pipeline is specifically designed to detect [5–8].

Scaffolds shorter than 1 kb are explicitly excluded from the final decontaminated assemblies. This threshold is necessary as short sequences generally lack sufficient k-mer diversity to enable robust compositional clustering and rarely contain enough complete open reading frames to support reliable taxonomic assignment [13,14]. Although some of these short scaffolds may correspond to genuine contaminant fragments, they more commonly represent assembly artifacts, often arising from repeat collapse or similar assembly graph ambiguities. By removing short scaffolds, the pipeline eliminates a pool of potential contaminants while predominantly sacrificing erroneous sequences.

Notably, we did not detect viral contaminants in the analyzed sponge genomes. Considering that our decontamination pipeline did not successfully recover viral sequences from the validation dataset, this absence most likely reflects methodological limitations rather than a true lack of viral signal. This may be attributable to incomplete representation of viral sequences in the databases utilized by our methodology, as well as to short viral scaffolds that lack sufficient discriminatory signal. Future iterations of the pipeline would benefit from incorporating dedicated viral discovery components, such as curated virus-specific reference databases or hallmark gene detection approaches.

A major challenge in contaminant detection is the potential for HGT to introduce false-positive contamination signals [5,6,8,36]. Because HGT candidates often harbor sequence features resembling microbial donors, they risk being erroneously targeted by decontamination algorithms [11,13,14]. Although we did not explicitly test the pipeline against a curated set of confirmed HGT events, the validation dataset included sponge chromosomal fragments likely harbouring ancestral HGTs and nevertheless yielded extremely high precision. This result suggests that the pipeline effectively distinguishes bona fide HGT-derived sequences from exogenous microbial contaminants.

The framework presented here is inherently adaptable and can be applied to other taxa. Although several decontamination tools have been developed [12–19], we did not perform direct benchmarking against alternative pipelines. The primary objective of this study was to design a reproducible, consensus-based workflow specifically tailored to non-model organisms and to apply it systematically across publicly available sponge genomes. Our goal was not to replace established tools but to provide a complementary strategy optimized for holobiont systems with limited representation in reference databases.

The taxonomic characterization of detected contaminants highlights the complex biological landscape of the sponge holobiont. Our analysis revealed a consistent dominance of specific bacterial phyla, most notably Pseudomonadota, Chloroflexota, and Poribacteria. This profile aligns closely with the established composition of high-microbial-abundance sponges, where these lineages are known to provide essential benefits, ranging from nitrogen cycling to the synthesis of secondary metabolites [5–8]. However, the diversity of contaminants varied significantly across the analyzed species, likely reflecting both the ecological characteristics of the host and the decontamination strategy used in the original study. Species such as *P*. *foliascens* and *X*. *muta* exhibited the highest contaminant diversity. In these assemblies, we identified not only dominant bacterial groups but also a significant amount of eukaryotic contaminants, including green algae and fungi. Notably, both algal and fungal lineages have been previously reported as sponge-associated endosymbionts, where they contribute to nutrient exchange and secondary metabolite production [4,6,37,38]. Archaeal lineages were also detected in several assemblies, with many of these taxa corresponding to ammonia-oxidizing species that play central roles in the nitrogen cycling of sponges [3,5,6,39]. Ultimately, these results suggest that contamination in sponge genomics may primarily be a reflection of the biological integration between the sponge and its microbial partners.

### Conclusion

Our results demonstrate that contamination in sponge genome assemblies is widespread yet can be effectively addressed through a consensus-based framework. By integrating compositional and taxonomy-based evidence, our decontamination pipeline achieves high accuracy while maintaining host genome completeness, even in the context of underrepresented and microbially complex systems. This study provides 30 curated sponge genome assemblies, thus offering a cleaner foundation for investigating the evolutionary and functional biology of sponges. Overall, our findings support the utility of a consensus-based decontamination strategy as a reliable framework for improving genome quality in non-model metazoans with complex microbial interactions.

## List of abbreviations

HGT: Horizontal Gene Transfer

CLR: Centered Log-ratio

PCA: Principal Component Analysis

## Declarations

### Ethics approval and consent to participate

Not applicable

### Consent for publication

Not applicable

### Availability of data and materials

The decontaminated versions of all 30 sponge genome assemblies generated in this study are publicly available on Zenodo (https://doi.org/10.5281/zenodo.19064896). Each assembly is provided in FASTA format, accompanied by MD5 checksums. For each analysed assembly, we also provide a detailed report including per-scaffold contamination statistics, taxonomic assignments of contaminant scaffolds, and BUSCO completeness metrics. The validation dataset results are also provided.

The code used to perform all analyses reported in this study is available on GitHub (https://github.com/bodulic/sponge_genome_decontamination_workflow) and Zenodo (https://doi.org/10.5281/zenodo.19139232).

The decontamination pipeline, documentation, and example commands are available under the MIT license on GitHub (https://github.com/bodulic/genome_decontamination) and Zenodo (https://doi.org/10.5281/zenodo.19068936).

### Authors’ contributions

Conceptualization, K.B., K.V.; data curation, K.B.; formal analysis, K.B.; investigation, K.B.; methodology, K.B.; project administration, K.V.; resources, K.V.; software, K.B.; supervision, K.V.; validation, K.B., K.V.; visualization, K.B.; writing of the original draft, K.B.; writing review and editing, K.V.

All authors read and approved the final manuscript.

## Acknowledgements

Not applicable

