## Supplementary information for "Hidden Contaminants in Sponge Genomes: Large-Scale Decontamination of 30 Public Assemblies"

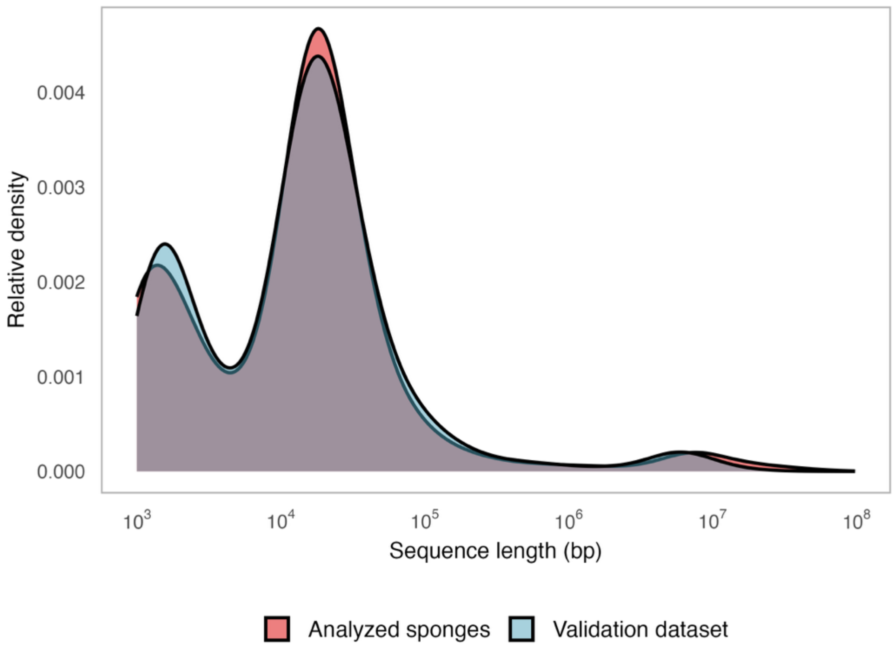


**Supplementary Figure 1.** Distribution of sequence length in the spike-in validation dataset and sponge genome assemblies analysed for contamination.

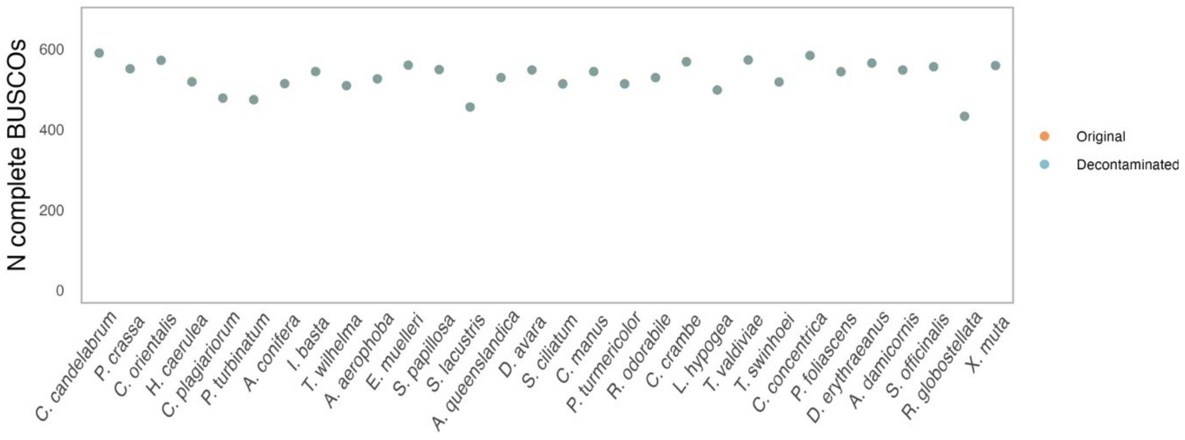


**Supplementary Figure 2**. Comparison of the number of complete BUSCO orthologs between the original and decontaminated assemblies (Metazoa lineage, maximum *n* = 672).

**Supplementary Table 1**. Default values of tunable parameters in the genome decontamination pipeline.

| Score / analysis | Parameter | Default value | Rationale |
|---|---|---|---|
| Compositional score | Maximum number of Gaussian finite mixture model clusters | 10 | The default maximum of 10 clusters was chosen to provide sufficient flexibility to capture compositional heterogeneity and potential contaminant signals while avoiding overfitting and excessive model complexity. Assemblies typically contain a limited number of major compositional groups (e.g., target genome, organelles, contaminants), and a maximum of 10 components provides ample capacity to capture this diversity |
| Compositional score | Chi-square significance level threshold for outlier cluster detection | 0.005 | The default significance level of 0.005 was chosen to provide a conservative criterion for identifying compositional outliers, reducing false positives arising from natural variation in k-mer composition while maintaining sensitivity to strong deviations indicative of contamination. A more stringent threshold than commonly used values (e.g., 0.05 or 0.01) is adopted as no formal correction for multiple comparisons is applied when evaluating cluster distances |
| Compositional score | GC content z-score threshold for outlier scaffold detection | 2.5 | The GC content z-score threshold of 2.5 was chosen to identify scaffolds with substantial deviations from the genome-wide GC distribution while avoiding spurious outlier calls due to intrinsic variation |
| Protein taxonomy score | Multiplying factor for the DIAMOND hit set weight comparison | 1.2 | A multiplying factor of 1.2 requires that non-target taxonomic signal exceeds the target signal by a meaningful margin, reducing sensitivity to minor fluctuations in alignment scores while retaining sensitivity to genuine contamination |
| Protein taxonomy score | Minimum non-target hit set coverage fraction of a scaffold required for contaminant classification | 0.2 | A minimum coverage fraction of 0.2 requires that non-target DIAMOND hit sets span a meaningful portion of the scaffold, reducing the influence of isolated matches while retaining sensitivity to contaminant scaffolds with a limited number of protein-coding regions |
| Nucleotide taxonomy score | Multiplying factor for the Kraken2 k-mer ratio comparison | 1.2 | This parameter follows the same rationale as the multiplying factor for the DIAMOND hit set weight comparison described above |
| Nucleotide taxonomy score | Minimum non-target Kraken2 k-mer fraction required for contaminant classification | 0.05 | A minimum absolute k-mer ratio of 0.05 requires that non-target k-mers constitute a significant portion of the scaffold, while ensuring sufficient sensitivity to contaminants not included in the NCBI core nucleotide database |
| Taxonomy assignment | Minimum Kraken2 k-mer fraction required to assign a taxon to a scaffold | 0.1 | These thresholds jointly ensure that a taxonomic assignment is supported by both sufficient absolute k-mer evidence and a strong relative signal compared to other taxa. The minimum k-mer fraction (0.1) requires that a meaningful portion of the scaffold supports the assignment, while the relative fraction (0.5) ensures that the assigned taxon represents the dominant signal, reducing spurious classifications |
| Taxonomy assignment | Minimum Kraken2 relative k-mer fraction required to assign a taxon to a scaffold | 0.5 | |
| Taxonomy assignment | Minimum DIAMOND hit set coverage fraction required to assign a taxon to a scaffold | 0.25 | These parameters follow the same rationale as nucleotide-based taxonomy assignment thresholds described above |
| Taxonomy assignment | Minimum DIAMOND hit set relative coverage fraction required to assign a taxon to a scaffold | 0.5 | |

**Supplementary Table 2**. Sponge genome assemblies included in the spike-in validation dataset.*

| Species | Class | Accession number** | Assembly size (Mb) | N scaffolds | N chromosomes | N50 (Mb) | L50 |
|---|---|---|---|---|---|---|---|
| *Agelas tubulata* | Demospongiae | GCA_964245335.1 | 280.8 | 56 | 23 | 12.1 | 7 |
| *Aiolochroia crassa* | Demospongiae | GCA_964263335.1 | 188.6 | 37 | 23 | 7.9 | 9 |
| *Aphrocallistes beatrix* | Hexactinellida | GCA_963281255.1 | 100.5 | 28 | 19 | 5.0 | 8 |
| *Aplysina cauliformis* | Demospongiae | GCA_965643675.1 | 197.7 | 50 | 22 | 8.2 | 8 |
| *Aplysina cavernicola* | Demospongiae | GCA_964659575.1 | 168.1 | 23 | 20 | 7.6 | 8 |
| *Bolosoma cyanae* | Hexactinellida | GCA_964258925.1 | 101.7 | 27 | 19 | 4.8 | 7 |
| *Chondrilla caribensis* | Demospongiae | GCA_964020155.1 | 137.1 | 46 | 25 | 5.3 | 12 |
| *Chondrosia reniformis* | Demospongiae | GCA_947172415.1 | 117.4 | 14 | 14 | 8.5 | 7 |
| *Geodia parva* | Demospongiae | GCA_964274975.1 | 113.3 | 48 | 21 | 4,9 | 8 |
| *Geodia phlegraei* | Demospongiae | GCA_964341465.1 | 112.6 | 42 | 21 | 5.0 | 7 |
| *Halichondria panicea* | Demospongiae | GCA_963675165.1 | 131.5 | 39 | 17 | 7.1 | 8 |
| *Oscarella lobularis* | Homoscleromorpha | GCA_947507565.1 | 65.2 | 21 | 20 | 3.5 | 8 |
| *Pachastrella ovisternata* | Demospongiae | GCA_964263345.1 | 166.8 | 26 | 21 | 7.2 | 7 |
| *Petrosia ficiformis* | Demospongiae | GCA_947044365.1 | 191.2 | 35 | 18 | 9.90 | 18 |
| *Phakellia ventilabrum* | Demospongiae | GCA_964276705.1 | 211.9 | 26 | 25 | 8.4 | 10 |
| *Spongia lamella* | Demospongiae | GCA_963930835.2 | 130.4 | 24 | 10 | 14.5 | 4 |
| *Weberella bursa* | Demospongiae | GCA_965644515.1 | 92.3 | 24 | 24 | 3.8 | 11 |
| *Xestospongia bergquistia* | Demospongiae | GCA_963965975.1 | 146.5 | 20 | 15 | 8.9 | 5 |

*The dataset also contained five mitochondrial sequences from the following sponges: *Bolosoma cyanae*, *Chondrosia reniformis*, *Petrosia ficiformis*, *Phakellia ventilabrum*, and *Weberella bursa*

**GenBank assembly accession number

**Supplementary Table 3**. Contaminant genome assemblies included in the spike-in validation dataset.*

| Species | Taxon | Class | Accession number** | Assembly size (Mb) | N scaffolds | N50 (Mb) | L50 |
|---|---|---|---|---|---|---|---|
| *Leisingera methylohalidivorans* | Bacteria | Alphaproteobacteria | GCA_000511355.1 | 4.7 | 3 | 4.1 | 1 |
| *Phaeobacter gallaeciensis* | Bacteria | Alphaproteobacteria | GCA_000511385.1 | 4.5 | 8 | 3.8 | 1 |
| *Pseudovibrio brasiliensis* | Bacteria | Alphaproteobacteria | GCA_018282095.1 | 6.0 | 6 | 4.8 | 1 |
| *Pseudovibrio denitrificans* | Bacteria | Alphaproteobacteria | GCA_047444875.1 | 5.9 | 3 | 4.9 | 1 |
| *Roseobacter litoralis* | Bacteria | Alphaproteobacteria | GCA_000154785.2 | 4.7 | 4 | 4.5 | 1 |
| *Ruegeria pomeroyi* | Bacteria | Alphaproteobacteria | GCA_000011965.2 | 4.6 | 2 | 4.1 | 1 |
| *Sulfitobacter pontiacus* | Bacteria | Alphaproteobacteria | GCA_040790665.1 | 3.6 | 5 | 3.0 | 1 |
| *Sulfitobacter rhodophyticola* | Bacteria | Alphaproteobacteria | GCA_041281685.2 | 4.4 | 6 | 4.0 | 1 |
| *Thalassospira xiamenensis* | Bacteria | Alphaproteobacteria | GCA_000300235.2 | 4.8 | 2 | 4.6 | 1 |
| *Alteromonas naphthalenivorans* | Bacteria | Gammaproteobacteria | GCA_000213655.1 | 5.0 | 1 | 5.0 | 1 |
| *Alteromonas stellipolaris* | Bacteria | Gammaproteobacteria | GCA_001562195.1 | 4.6 | 1 | 4.6 | 1 |
| *Marinobacter nauticus* | Bacteria | Gammaproteobacteria | GCA_000284615.1 | 4.0 | 1 | 4.0 | 1 |
| *Pseudomonas putida* | Bacteria | Gammaproteobacteria | GCA_000412675.1 | 6.2 | 1 | 6.2 | 1 |
| *Vibrio vulnificus* | Bacteria | Gammaproteobacteria | GCA_002224265.1 | 5.0 | 2 | 3.3 | 1 |
| *Desulfobacter hydrogenophilus* | Bacteria | Deltaproteobacteria | GCA_004319545.1 | 5.3 | 3 | 5.2 | 1 |
| *Desulfobacter postgatei* | Bacteria | Deltaproteobacteria | GCA_000233695.3 | 4.0 | 1 | 4.0 | 1 |
| *Desulfobulbus propionicus* | Bacteria | Deltaproteobacteria | GCA_000186885.1 | 3.9 | 1 | 3.9 | 1 |
| *Desulfovibrio mangrovi* | Bacteria | Deltaproteobacteria | GCA_026230175.1 | 3.9 | 1 | 3.9 | 1 |
| *Geobacter sulfurreducens* | Bacteria | Deltaproteobacteria | GCA_000007985.2 | 3.8 | 1 | 3.8 | 1 |
| *Bacillus cabrialesii* | Bacteria | Bacilli | GCA_004124315.2 | 4.1 | 1 | 4.1 | 1 |
| *Bacillus velezensis* | Bacteria | Bacilli | GCA_000015785.2 | 3.9 | 1 | 3.9 | 1 |
| *Oceanobacillus iheyensis* | Bacteria | Bacilli | GCA_000011245.1 | 3.6 | 1 | 3.6 | 1 |
| *Virgibacillus natechei* | Bacteria | Bacilli | GCA_026013645.1 | 3.9 | 1 | 3.9 | 1 |
| *Virgibacillus pantothenticus* | Bacteria | Bacilli | GCA_018075365.1 | 4.8 | 1 | 4.8 | 1 |
| *Occallatibacter riparius* | Bacteria | Acidobacteriia | GCA_025264625.1 | 6.8 | 1 | 6.8 | 1 |
| *Micrococcus luteus* | Bacteria | Actinobacteria | GCA_900475555.1 | 2.5 | 1 | 2.5 | 1 |
| *Micrococcus porci* | Bacteria | Actinobacteria | GCA_020097155.1 | 2.6 | 1 | 2.6 | 1 |
| *Streptomyces collinus* | Bacteria | Actinobacteria | GCA_031348265.1 | 8.9 | 1 | 8.9 | 1 |
| *Leptolyngbya boryana* | Bacteria | Cyanophyceae | GCA_002142475.1 | 6.8 | 4 | 6.2 | 1 |
| *Prochlorococcus marinus* | Bacteria | Cyanophyceae | GCA_000015665.1 | 1.7 | 1 | 1.7 | 1 |
| *Synechococcus elongatus* | Bacteria | Cyanophyceae | GCA_022984195.1 | 2.7 | 3 | 2.7 | 1 |
| *Niabella soli* | Bacteria | Chitinophagia | GCA_000243115.3 | 4.7 | 1 | 4.7 | 1 |
| *Echinicola vietnamensis* | Bacteria | Cytophagia | GCA_000325705.1 | 5.6 | 1 | 5.6 | 1 |
| *Dehalococcoides mccartyi* | Bacteria | Dehalococcoidia | GCA_000011905.1 | 1.5 | 1 | 1.5 | 1 |
| *Dehalogenimonas formicexedens* | Bacteria | Dehalococcoidia | GCA_001953175.1 | 2.1 | 1 | 2.1 | 1 |
| *Formosa sediminum* | Bacteria | Flavobacteria | GCA_007197735.1 | 3.9 | 1 | 3.9 | 1 |
| *Halobacterium salinarum* | Bacteria | Halobacteria | GCA_004799605.1 | 2.4 | 3 | 2.2 | 1 |
| *Halorubrum ruber* | Bacteria | Halobacteria | GCA_018228765.1 | 3.0 | 1 | 3.0 | 1 |
| *Nitrospira japonica* | Bacteria | Nitrospira | GCA_900169565.1 | 4.1 | 1 | 4.1 | 1 |
| *Nitrospira moscoviensis* | Bacteria | Nitrospira | GCA_001273775.1 | 4.6 | 1 | 4.6 | 1 |
| *Haloferula helveola* | Bacteria | Planctomycetia | GCA_037076345.1 | 5.7 | 1 | 5.7 | 1 |
| *Planctomyces* sp. | Bacteria | Planctomycetia | GCA_001610835.1 | 8.4 | 1 | 8.4 | 1 |
| *Rhodopirellula baltica* | Bacteria | Planctomycetia | GCA_000196115.1 | 7.1 | 1 | 7.1 | 1 |
| *Paludibaculum fermentans* | Bacteria | Sphingobacteriia | GCA_015277775.1 | 9.5 | 2 | 9.5 | 1 |
| *Verrucomicrobium spinosum* | Bacteria | Verrucomicrobiae | GCA_000172155.1 | 8.2 | 1 | 8.2 | 1 |
| *Methanobrevibacter smithii* | Archaea | Methanobacteria | GCA_000016525.1 | 1.9 | 1 | 1.9 | 1 |

| | | | | | | | |
|---|---|---|---|---|---|---|---|
| *Nitrosopelagicus brevis* | Archaea | Nitrososphaeria | GCA_000812185.1 | 1.2 | 1 | 1.2 | 1 |
| *Nitrosotenuis cloacae* | Archaea | Nitrososphaeria | GCA_000955905.3 | 1.6 | 1 | 1.6 | 1 |
| *Nitrosopumilus maritimus* | Archaea | Nitrososphaeria | GCA_000018465.1 | 1.6 | 1 | 1.6 | 1 |
| *Cryptosporidium parvum* | Protozoa | Conoidasida | GCA_000165345.1 | 9.1 | 8 | 1.1 | 4 |
| *Dictyostelium discoideum* | Protozoa | Dictyostelia | GCA_000004695.1 | 34.1 | 50 | 5.5 | 3 |
| *Acanthamoeba castellanii* | Protozoa | Discosea | GCA_000313135.1 | 42.0 | 384 | 0.3 | 26 |
| *Capsaspora owczarzaki* | Protozoa | Filasterea | GCA_000151315.2 | 28.0 | 84 | 1.6 | 6 |
| *Paramecium tetraurelia* | Protozoa | Oligohymenophorea | GCA_000165425.1 | 72.1 | 697 | 0.4 | 64 |
| *Tetrahymena thermophila* | Protozoa | Oligohymenophorea | GCA_000189635.1 | 103.0 | 1150 | 0.5 | 59 |
| *Phaeodactylum tricornutum* | Algae | Bacillariophyceae | GCA_000150955.2 | 27.5 | 88 | 1.0 | 11 |
| *Thalassiosira pseudonana* | Algae | Bacillariophyceae | GCA_000149405.2 | 32.4 | 64 | 2.0 | 7 |
| *Ostreococcus tauri* | Algae | Mamiellophyceae | GCA_000214015.2 | 12.9 | 20 | 0.8 | 7 |
| *Chlorella sorokiniana* | Algae | Trebouxiophyceae | GCA_025917655.1 | 38.6 | 13 | 3.0 | 5 |
| *Aspergillus sydowii* | Fungi | Eurotiomycetes | GCA_001890705.1 | 34.4 | 97 | 2.3 | 5 |
| *Malassezia globosa* | Fungi | Malasseziomycetes | GCA_000181695.2 | 9.0 | 62 | 0.7 | 5 |
| *Candida parapsilosis* | Fungi | Saccharomycetes | GCA_000182765.2 | 13.0 | 8 | 2.1 | 3 |
| *Debaryomyces hansenii* | Fungi | Saccharomycetes | GCA_000006445.2 | 12.2 | 7 | 2.0 | 3 |
| *Spizellomyces punctatus* | Fungi | Spizellomycetes | GCA_000182565.2 | 24.1 | 38 | 1.5 | 7 |
| Alteromonas phage | Viruses | Caudoviricetes | GCA_010706295.1 | 0.09 | 1 | 0.09 | 1 |
| Ruegeria phage | Viruses | Caudoviricetes | GCA_000925815.1 | 0.06 | 1 | 0.06 | 1 |
| Vibrio phage VP16T | Viruses | Caudoviricetes | GCA_004216455.1 | 0.05 | 1 | 0.05 | 1 |
| Vibrio phage CTXphi | Viruses | Faserviricetes | GCA_000893195.1 | 0.01 | 1 | 0.01 | 1 |
| Chaetoceros socialis RNA virus 01 | Viruses | Pisoniviricetes | GCA_000882095.1 | 0.01 | 1 | 0.01 | 1 |
| Heterosigma akashiwo RNA virus | Viruses | Pisoniviricetes | GCA_000863545.1 | 0.01 | 1 | 0.01 | 1 |
| Acanthamoeba polyphaga mimivirus | Viruses | Megaviricetes | GCA_000888735.1 | 1.2 | 1 | 1.2 | 1 |

*The dataset also contained five plasmid sequences from the following bacteria: *Desulfobacter hydrogenophilus* (RefSeq NZ_CP036314.1), *Paludibaculum fermentans* (RefSeq NZ_CP063850.1), *Pseudovibrio brasiliensis* (RefSeq NZ_CP074127.1), *Synechococcus elongatus* (RefSeq NZ_CP085786.1), and *Thalassospira xiamenensis* (RefSeq NZ_CP004389.1)

**GenBank assembly accession number

**Supplementary Table 4**. Sponge genome assemblies included in the contamination analysis.

| Species | Class | Accession number* | Assembly size (Mb) | N scaffolds | N50 (Mb) | L50 |
|---|---|---|---|---|---|---|
| *Agelas conifera* | Demospongiae | GCA_965122385.1 | 338.6 | 2617 | 9.5 | 9 |
| *Amphimedon queenslandica* | Demospongiae | GCA_016292275.1 | 167.7 | 3871 | 0.95 | 49 |
| *Aplysina aerophoba* | Demospongiae | GCA_949841015.1 | 158 | 158 | 6.6 | 10 |
| *Axinella damicornis* | Demospongiae | GCA_963931865.1 | 234.9 | 663 | 15.7 | 6 |
| *Callyspongia manus* | Demospongiae | GCA_965112345.1 | 486.3 | 339 | 32.0 | 7 |
| *Cliona orientalis* | Demospongiae | GCA_963930775.1 | 217.2 | 307 | 10.0 | 8 |
| *Corbitella plagiariorum* | Hexactinellida | GCA_965278765.1 | 111.8 | 873 | 4.2 | 11 |
| *Corticium candelabrum* | Homoscleromorpha | GCA_963422355.1 | 185.5 | 115 | 8.5 | 10 |
| *Crambe crambe* | Demospongiae | GCA_963924555.1 | 143.2 | 124 | 7.7 | 8 |
| *Cymbastela concentrica* | Demospongiae | GCA_965112925.1 | 216.4 | 149 | 8.4 | 11 |
| *Diacarnus erythraeanus* | Demospongiae | GCA_964016965.1 | 140.9 | 145 | 7.5 | 8 |
| *Dysidea avara* | Demospongiae | GCA_963678975.2 | 550.7 | 108 | 40.5 | 6 |
| *Ephydatia muelleri* | Demospongiae | GCA_049114765.1 | 230.3 | 165 | 8.90 | 9 |
| *Halisarca caerulea* | Demospongiae | GCA_963170055.1 | 195.7 | 488 | 2.8 | 19 |
| *Ianthella basta* | Demospongiae | GCA_964019415.1 | 209.3 | 134 | 8.3 | 9 |
| *Lycopodina hypogea* | Demospongiae | GCA_963969325.1 | 235.1 | 199 | 16.0 | 7 |
| *Penares turmericolor* | Demospongiae | GCA_965152365.1 | 130.7 | 173 | 5.6 | 9 |
| *Petrosia crassa* | Demospongiae | GCA_965785925.1 | 226.9 | 112 | 9.2 | 4 |
| *Phyllospongia foliascens* | Demospongiae | GCA_964212625.1 | 403.0 | 1807 | 23.8 | 6 |
| *Pleroma turbinatum* | Demospongiae | GCA_965278755.2 | 131.2 | 1650 | 3.7 | 8 |
| *Rhabdastrella globostellata* | Demospongiae | GCA_964261875.1 | 155.6 | 328 | 6.5 | 6 |
| *Rhopaloeides odorabile* | Demospongiae | GCA_964237215.1 | 291.6 | 374 | 16.4 | 7 |
| *Spheciospongia papillosa* | Demospongiae | GCA_965235015.2 | 251.4 | 1837 | 10.8 | 9 |
| *Spongia officinalis* | Demospongiae | GCA_964213935.1 | 466.2 | 105 | 28.5 | 6 |
| *Spongilla lacustris* | Demospongiae | GCA_949361645.1 | 248.7 | 70 | 10.1 | 8 |
| *Sycon ciliatum* | Calcarea | GCF_964019385.1 | 539.6 | 618 | 39.0 | 5 |
| *Tethya wilhelma* | Demospongiae | GCA_964030475.1 | 126.2 | 557 | 6.7 | 4 |
| *Thenea valdiviae* | Demospongiae | GCA_964340815.1 | 173.8 | 230 | 8.1 | 8 |
| *Theonella swinhoei* | Demospongiae | GCA_963970385.1 | 144.1 | 168 | 6.1 | 8 |
| *Xestospongia muta* | Demospongiae | GCA_963693285.1 | 158.5 | 298 | 10.6 | 4 |

*RefSeq / GenBank assembly accession number

**Supplementary Table 5**. Contamination statistics for the analysed sponge genome assemblies.

| | Contamination statistics | | Contaminant taxonomy | | | | | |
|---|---|---|---|---|---|---|---|---|
| **Species** | **Contaminant scaffold N (%)** | **Contaminant length (Mb, %)** | **Bacteria N (%)** | **Archaea N (%)** | **Protozoa N (%)** | **Algae N (%)** | **Fungi N (%)** | **Other* N (%)** |
| *Agelas conifera* | 15 (0.6%) | 0.34 (1.00%) | 11 (0.4%) | 0 (0.0%) | 0 (0.0%) | 0 (0.0%) | 0 (0.0%) | 4 (0.2%) |
| *Amphimedon queenslandica* | 191 (4.9%) | 2.05 (1.22%) | 150 (3.9%) | 0 (0.0%) | 3 (0.1%) | 2 (0.1%) | 4 (0.1%) | 32 (0.8%) |
| *Aplysina aerophoba* | 7 (4.4%) | 0.27 (0.17%) | 1 (0.6%) | 0 (0.0%) | 1 (0.6%) | 1 (0.6%) | 1 (0.6%) | 3 (1.9%) |
| *Axinella damicornis* | 239 (36.0%) | 5.79 (2.47%) | 0 (0.0%) | 1 (0.2%) | 18 (2.7%) | 168 (25.3%) | 46 (6.9%) | 6 (0.9%) |
| *Callyspongia manus* | 31 (9.1%) | 0.53 (0.11%) | 22 (6.5%) | 2 (0.6%) | 0 (0.0%) | 4 (1.2%) | 0 (0.0%) | 3 (0.9%) |
| *Cliona orientalis* | 1 (0.3%) | 0.02 (0.01%) | 0 (0.0%) | 0 (0.0%) | 0 (0.0%) | 1 (0.3%) | 0 (0.0%) | 0 (0.0%) |
| *Corbitella plagiariorum* | 4 (0.5%) | 0.27 (0.24%) | 2 (0.2%) | 0 (0.0%) | 1 (0.1%) | 0 (0.0%) | 0 (0.0%) | 1 (0.1%) |
| *Corticium candelabrum* | 0 (0.0%) | 0.00 (0.00%) | 0 (0.0%) | 0 (0.0%) | 0 (0.0%) | 0 (0.0%) | 0 (0.0%) | 0 (0.0%) |
| *Crambe crambe* | 15 (12.1%) | 0.24 (0.17%) | 10 (8.1%) | 2 (1.6%) | 0 (0.0%) | 0 (0.0%) | 1 (0.8%) | 2 (1.6%) |
| *Cymbastela concentrica* | 26 (17.4%) | 0.92 (0.43%) | 2 (1.3%) | 0 (0.0%) | 0 (0.0%) | 16 (10.7%) | 2 (1.3%) | 6 (4.0%) |
| *Diacarnus erythraeanus* | 50 (34.5%) | 0.85 (0.60%) | 16 (11.0%) | 2 (1.4%) | 1 (0.7%) | 25 (17.2%) | 1 (0.7%) | 5 (3.4%) |
| *Dysidea avara* | 8 (7.4%) | 0.13 (0.02%) | 5 (4.6%) | 1 (0.9%) | 0 (0.0%) | 1 (0.9%) | 0 (0.0%) | 1 (0.9%) |
| *Ephydatia muelleri* | 8 (4.8%) | 0.19 (0.08%) | 5 (3.0%) | 0 (0.0%) | 0 (0.0%) | 1 (0.6%) | 1 (0.6%) | 1 (0.6%) |
| *Halisarca caerulea* | 2 (0.4%) | 0.39 (0.18%) | 0 (0.0%) | 0 (0.0%) | 0 (0.0%) | 2 (0.4%) | 0 (0.0%) | 0 (0.0%) |
| *Ianthella basta* | 2 (1.5%) | 0.08 (0.04%) | 1 (0.7%) | 0 (0.0%) | 0 (0.0%) | 0 (0.0%) | 0 (0.0%) | 1 (0.7%) |
| *Lycopodina hypogea* | 29 (14.6%) | 0.29 (0.12%) | 25 (12.6%) | 1 (0.5%) | 0 (0.0%) | 1 (0.5%) | 0 (0.0%) | 2 (1.0%) |
| *Penares turmericolor* | 18 (10.4%) | 0.34 (0.26%) | 10 (5.8%) | 6 (3.5%) | 0 (0.0%) | 1 (0.6%) | 0 (0.0%) | 1 (0.6%) |
| *Petrosia crassa* | 0 (0.0%) | 0.00 (0.00%) | 0 (0.0%) | 0 (0.0%) | 0 (0.0%) | 0 (0.0%) | 0 (0.0%) | 0 (0.0%) |
| *Phyllospongia foliascens* | 562 (31.1%) | 14.15 (3.51%) | 188 (10.4%) | 2 (0.1%) | 35 (1.9%) | 195 (10.8%) | 96 (5.3%) | 46 (2.5%) |
| *Pleroma turbinatum* | 9 (0.5%) | 0.40 (0.30%) | 9 (0.5%) | 0 (0.0%) | 0 (0.0%) | 0 (0.0%) | 0 (0.0%) | 0 (0.0%) |
| *Rhabdastrella globostellata* | 156 (47.6%) | 0.54 (0.34%) | 126 (38.4%) | 0 (0.0%) | 1 (0.3%) | 0 (0.0%) | 1 (0.3%) | 28 (8.5%) |
| *Rhopaloeides odorabile* | 43 (11.5%) | 0.68 (0.23%) | 24 (6.4%) | 1 (0.3%) | 0 (0.0%) | 0 (0.0%) | 0 (0.0%) | 18 (4.8%) |
| *Spheciospongia papillosa* | 96 (5.2%) | 2.32 (0.92%) | 72 (3.9%) | 3 (0.2%) | 2 (0.1%) | 11 (0.6%) | 4 (0.2%) | 4 (0.2%) |
| *Spongia officinalis* | 40 (38.1%) | 0.63 (0.14%) | 24 (22.9%) | 1 (1.0%) | 1 (1.0%) | 3 (2.9%) | 0 (0.0%) | 11 (10.5%) |
| *Spongilla lacustris* | 4 (5.7%) | 0.05 (0.02%) | 2 (2.9%) | 0 (0.0%) | 1 (1.4%) | 0 (0.0%) | 0 (0.0%) | 1 (1.4%) |
| *Sycon ciliatum* | 48 (7.8%) | 1.27 (0.24%) | 37 (6.0%) | 0 (0.0%) | 1 (0.2%) | 6 (1.0%) | 1 (0.2%) | 3 (0.5%) |
| *Tethya wilhelma* | 13 (2.3%) | 0.35 (0.27%) | 2 (0.4%) | 1 (0.2%) | 1 (0.2%) | 0 (0.0%) | 1 (0.2%) | 8 (1.4%) |
| *Thenea valdiviae* | 34 (14.8%) | 0.76 (0.44%) | 22 (9.6%) | 3 (1.3%) | 1 (0.4%) | 1 (0.4%) | 1 (0.4%) | 6 (2.6%) |
| *Theonella swinhoei* | 26 (15.5%) | 0.64 (0.44%) | 2 (1.2%) | 0 (0.0%) | 0 (0.0%) | 16 (9.5%) | 6 (3.6%) | 2 (1.2%) |
| *Xestospongia muta* | 200 (67.1%) | 0.42 (0.27%) | 146 (49.0%) | 4 (1.3%) | 0 (0.0%) | 1 (0.3%) | 1 (0.3%) | 48 (16.1%) |

*Corresponds to scaffolds with unresolved phylum-level taxonomy
